# Novel CAF-identifiers via transcriptomic analysis in oral cancer patients

**DOI:** 10.1101/2023.01.10.523511

**Authors:** Nehanjali Dwivedi, Nidhi Shukla, Manjula Das, Sujan K Dhar

**Affiliations:** Molecular Immunology, Mazumdar Shaw Medical foundation, 8^th^ Floor MSMC, Narayana Health City, Bommasandra, Hosur Road, Bangalore, Karnataka – 560099, India; Computational Biology, Mazumdar Shaw Medical foundation, 8^th^ Floor MSMC, Narayana Health City, Bommasandra, Hosur Road, Bangalore, Karnataka – 560099, India; Manipal, MAHE, 576104

**Keywords:** Oral cancer, Cancer Associated Fibroblasts, Patient derived cultures, CAF Markers, Single-cell sequence, RNA-seq

## Abstract

**Background:** Cancer-associated fibroblasts (CAFs), a prominent component of the tumor microenvironment, plays an important role in tumor development, invasion, and drug resistance. The expression of distinct “CAF markers,” which separates CAFs from normal fibroblasts and epithelial cells, have traditionally been used to identify them. These commonly used CAF markers have been reported to differ greatly across microenvironmental subpopulations even within a cancer site.

**Methods:** Using an unbiased data analysis approach utilizing publicly available and in-house gene expression data from patient derived novel CAF cells, we identified a collection of markers in oral cancer to distinguish CAF populations from tumor epithelia and normal oral fibroblasts.

**Results:** COL1A1, SPARC, COL1A2, COL3A1 and TIMP-1 were identified as potential markers which can be utilized to differentiate cancer associated fibroblast from all other cell types including normal fibroblasts in oral cancer.

## Background

Tumor microenvironment (TME) consists of dynamic cell populations such as fibroblasts, immune cells, inflammatory cells, adipocytes, endothelial cells, mesenchymal cells, and extracellular matrix (ECM) (1–3). Cross-talks between CAF and TME signal the tumor to survive and advance (4,5). During cancer development, there is a prominent expansion of quiescent fibroblasts residing in the host tissue in response to the developing neoplasm (6). CAFs have been recently defined decisively as cells negative for epithelial (EpCAM), endothelial (CD31) and leukocyte (CD45) markers with an elongated morphology and lacking the mutations found within cancer cells (7–10).

Multiple markers have been described to define CAFs for years. For example, Fibroblast Activation Protein (FAP) and Smooth Muscle Actin (SMA/ACTA2) have been employed as a marker of activated cancer-associated fibroblasts (8,11,12) respectively in colorectal and breast cancer because of their high expression in the tumor stroma. However, a number of investigations have revealed that epithelial cells undergoing epithelial-mesenchymal transition (EMT) also exhibit higher levels of FAP (13), while alpha SMA displays fluctuating expression amongst various CAF subtypes (14,15). Additionally, the intracellular location of alpha SMA makes it unsuitable for flow-sorting CAF populations for subsequent functional experiments. Although neither PDGFRα nor PDGFRβ are significantly upregulated in CAF populations, due to a more stable expression which is not sensitive to environmental variables like hypoxia, both have been considered as CAF markers in breast cancer (16). Similarly, Vimentin’s ubiquitous expression in the whole fibroblast population as well as multiple other cell types, including macrophages, adipocytes, and the cells undergoing EMT severely limits its utility as a CAF-specific marker (17–19). Expression of FSP1 (20), Transgelin (TAGLN) and Periostin (POSTN) (21) in CAFs vary among subtypes as reported in colorectal cancer (15). Podoplanin (PDPN), a membrane-bound marker that has been utilised to identify pro-tumorigenic fibroblast subpopulations lacks specificity since it is also expressed in epithelial tumor cells and inflammatory macrophages (22,23). Integrin 11-1 (ITGA11) has been identified to be upregulated in CAFs from non-small cell lung cancer (24,25), and has been demonstrated to be present in a variety of tumor cell lines, However, it is sensitive to factors like hypoxia (26), and TGF-signalling (27), which play a role as both an inducer and antagonist of specific CAF subtypes (28). Though described a few years back, Microfibril Associated Protein 5 (MFAP5) and Collagen Type XI Alpha I Chain (COL11A1) have not yet found popularity among the users. Further research is therefore necessary to characterise these markers’ activity and expression in various CAF subtypes (29,30).

Various factors secreted by CAF, have been implicated in promoting oral cancer invasion and/or proliferation, including IL-1β (interleukin 1β), activin A, HGF (hepatocyte growth factor) and EREG (epiregulin), which has additionally been suggested to play an autocrine role in CAF activation (31). CAF markers, reported so far in oral cancer has been summarized in Supplementary Table 1. With the development of single cell transcriptomic sequencing (scRNA-seq), it is now known that not all CAF subpopulations exhibit high expression of the most widely used marker – ACTA2, and that not all ACTA2-positve CAF are myofibroblasts. The molecular characteristics of oral cancer may also potentially affect the fibroblast phenotype. Puram et al. used scRNA-seq, and identified two primary fibroblast subgroups called “myofibroblasts” and “CAF.” Transcriptomic study of CAF cultures from oral malignancies (OSCC) by Patel et al. (32) resulted in the identification of two major subgroups based on ACTA2 expression: CAF1 (ACTA2-low) and CAF2 (ACTA2-high). CAF1 was linked to enhanced cancer cell proliferation but decreased self-renewal resembling oral stem cells (oral-SLCCs). On the other hand, CAF2 had a negative correlation with tumour cell proliferation and a positive correlation with the frequency of oral-SLCCs. Hassona et al. (33) reported that the cultured CAF from the genetically unstable OSCC (GU-OSCC) were considerably more senescent when compared to the genetically stable OSCC (GS-OSCC; with wild-type p53). They discovered that (myo)fibroblast activation, elevated production of TGFbeta-1 and TGFbeta-2, and high levels of reactive oxygen species (ROS) are all related with malignant keratinocytes from GU-OSCC. Senescent fibroblasts on the other hand, frequently express ACTA2, are contractile, and accelerate tumour growth (34,35), but cannot deposit ECM, as opposed to myofibroblasts (34,36). Interestingly, the production of intracellular ROS, specifically NADPH oxidase 4 (NOX4), is crucial to the development of CAF as well as myofibroblast (37,38).

Multiple methods, as described in Supplementary Table 2, have been employed to arrive at a set of CAF markers to distinguish them from normal fibroblast and epithelial cells. Since the development of sequencing technologies, availability of multi-omics data has transformed medicine and biology by allowing integrated system-level techniques to be developed (39). Researchers may compare, compute, and analyse the expression patterns of multiple genes across samples and cell types using gene expression analyses methods such as microarray, mRNA sequencing, and single cell sequencing, which are more robust as compared to the gold standard immunohistochemistry (IHC) methodology. Using highly sensitive single cell sequencing approach, Puram et.al, described two sets of CAFs – CAF1 and CAF2 having differential expression of immediate early response genes (e.g. JUN, FOS), mesenchymal markers (e.g. VIM, THY1), ligands and receptors (e.g. FGF7, TGFBR2/3), and ECM proteins (e.g. MMP11, CAV1) (40). However, experimental validation of the identified markers on pure primary cultures becomes indispensable to support the omics data in localization studies. Thus, it is important to consider all these aspects to analyse the markers before reporting. Due to different activation mechanisms and molecular characteristics (41,42), CAFs have greater functional diversity in various cancers (43–45). Two distinct CAF subtypes with different intratumoral localization have been recognized in pancreatic ductal adenocarcinoma (PDAC). Inflammatory CAFs (iCAFs), which secrete higher levels of Interleukin-6 (IL-6) along with other cytokines, are situated distantly from the tumor cells, whereas myofibroblastic CAFs (myCAFs) express alpha SMA and interact directly with neoplastic cells (46). Similar reports have been identified in breast cancer, where, Friedman et al. identified two major subpopulations of CAFs, namely pCAFs and sCAFs, categorised by the expression levels of podoplanin and S100A, respectively. During cancer progression, the molecular makeup and functionality of CAFs vary which is a predictor of the patient prognosis (47). Similarly in colorectal cancer (CRC), two subtypes of CAFs were identified: CAF-A and CAF-B, where, one category of CAFs displayed myCAFs markers i.e. ACTA2, TAGLN and PDGFA, whereas these markers were downregulated in the second CAF subset, which majorly expressed markers like DCN, MMP2, and COL1A2(48). Thus, unlike the tumor epithelia, functionally diverse CAFs from different cancers are difficult to define with one or two consistent markers.

Since there are no single definitive marker for CAFs and many positive markers utilised lack of specificity, negative selection is commonly used to filter out the cell types that are frequently present in tumor microenvironment. In order to distinguish between epithelial and smooth muscle cells, markers like epithelium cell adhesion molecule (EpCAM) and Smoothelin (SMTN) are frequently utilised (18,49,50). Leukocytes and endothelial cells are typically excluded using CD45, CD34, and CD11b (9,51).

As evident, it is unlikely to have a single or a single set of CAF markers for all cancer type. Without an -omic approach it is also not possible to have a common marker for various CAF types for any one cancer type.

The present study therefore employed an integrative analysis of publicly available single-cell and bulk gene expression data and in-house RNA-seq data to propose reliable CAF markers, which can then be validated on patient derived cells by various expression analysis techniques.

## Methods

The overall study workflow begins with sequencing of patient derived cells, data collection, and ends with final candidate markers using various levels of filters like (i) comparison of CAF and normal fibroblast transcriptome (ii) normal vs. tumor transcriptome from a public database, and a final filter of (iii) biological significance (Fig. 1).

**Figure 1.**
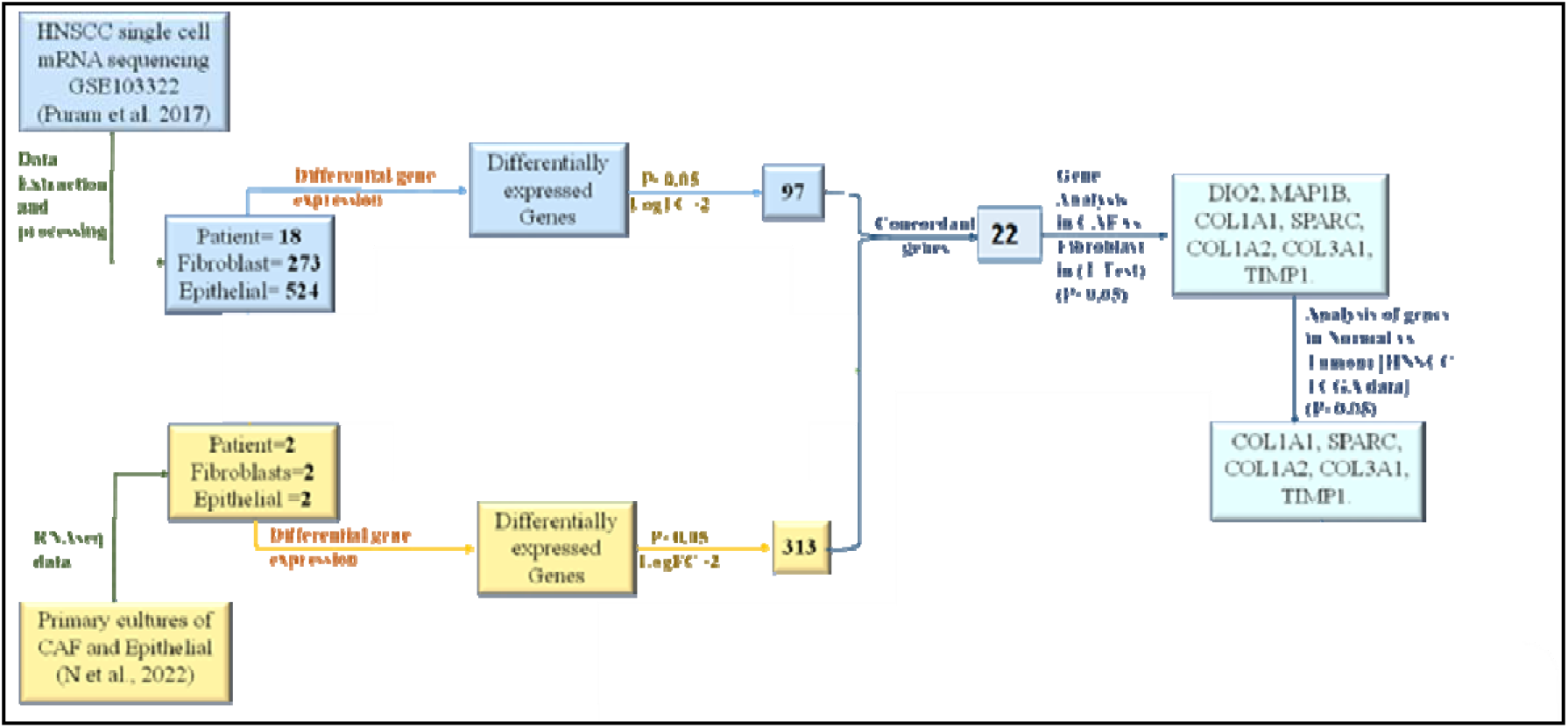
Workflow for identification of CAF-marker candidates. Data analysis was performed from public databases and in-house RNA-seq data to arrive at a concordant set of differentially expressed genes between epithelial and CAFs which were further selected over normal fibroblasts (Lai et.al, 2019), and adjacent normal in TCGA_HNSC. Markers involved in unique pathway were chosen using STRING database by interacting partner analysis. The chosen markers were then validated experimentally in patient derived CAF cells using qPCR, FACS and fluorescence microscopy.

### mRNA Sequencing of Patient Derived Cell Lines

Sequencing of mRNA extracted from MhCT08/12-E and MhCT08/12-F wa using illumina HiSeq series X sequencer using 150 bp paired-end chemistry. Reads obtained from the sequencer were quality checked and filtered using fastp (52) and subsequently aligned to GRCh38 reference assembly using STAR (53) pipeline.

### Acquisition of single cell sequencing data

Search with the terms (((HNSCC OR “ORAL CANCER” OR HNSCC) AND (single-cell OR “single cell”)) with organism filter for home sapiens in Gene Expression Omnibus (54) yielded three series with accession codes - GSE103322, GSE135975 and GSE163872. Out of these, GSE103322 (40) single-cell gene expression processed data and GSE135975 (55) RNA-seq FPKM (fragments per kilobase of transcript per million mapped reads) values were retrieved from GEO portal. Data from GSE163872 was not considered since the clinical details of the samples were not available.

### Data Extraction and Processing

GSE103322 data consisted of sequenced transcriptomes of roughly 6,000 single cells from 18 HNSCC patients containing 273 and 524 non-lymphatic fibroblast and epithelial cells. In parallel, RNA seq data from two novel autologous pairs of CAFs and epithelial cells were also analysed.

### Selection of putative CAF markers

Over-expressed genes in fibroblasts compared to epithelials in dataset GSE103322 were enumerated using the limma R package. Overexpressed genes in fibroblasts with log Fold Change value ≥ 2 and Benjamini-Hochberg adjusted p-value ≤ 0.05 from the in-house cell line sequencing data and from dataset GSE103322 were combined to identify genes concordant in both differential expression sets. The concordant genes were further filtered for differential expression between 8 patient-derived CAFs and 3 normal fibroblasts (n dataset GSE135975) using t-test to select genes over-expressed CAFs. Expression of these shortlisted genes were assessed in the TCGA Head and Neck Squamous Cell carcinoma dataset using RNA-seq expression values acquired from Broad Institute Firehose portal(56) to obtain set of genes overexpressed in tumor compared to its normal counterpart.

## Results

### Identification of differentially expressed genes between CAF and epithelial cells

Differential gene expression analysis of established CAF MhCT08/12-F and epithelial MhCT08/12-E patient derived cell lines revealed 313 significantly upregulated genes (Fig. 2a), whereas single-cell sequencing data from non-lymphatic fibroblast and epithelial cells obtained from GSE103322 (40) revealed 97 genes that were up-regulated in CAFs as compared to epithelial cells (Fig. 2b). 22 genes were concordant between the two differential expression subsets.

**Figure 2.**
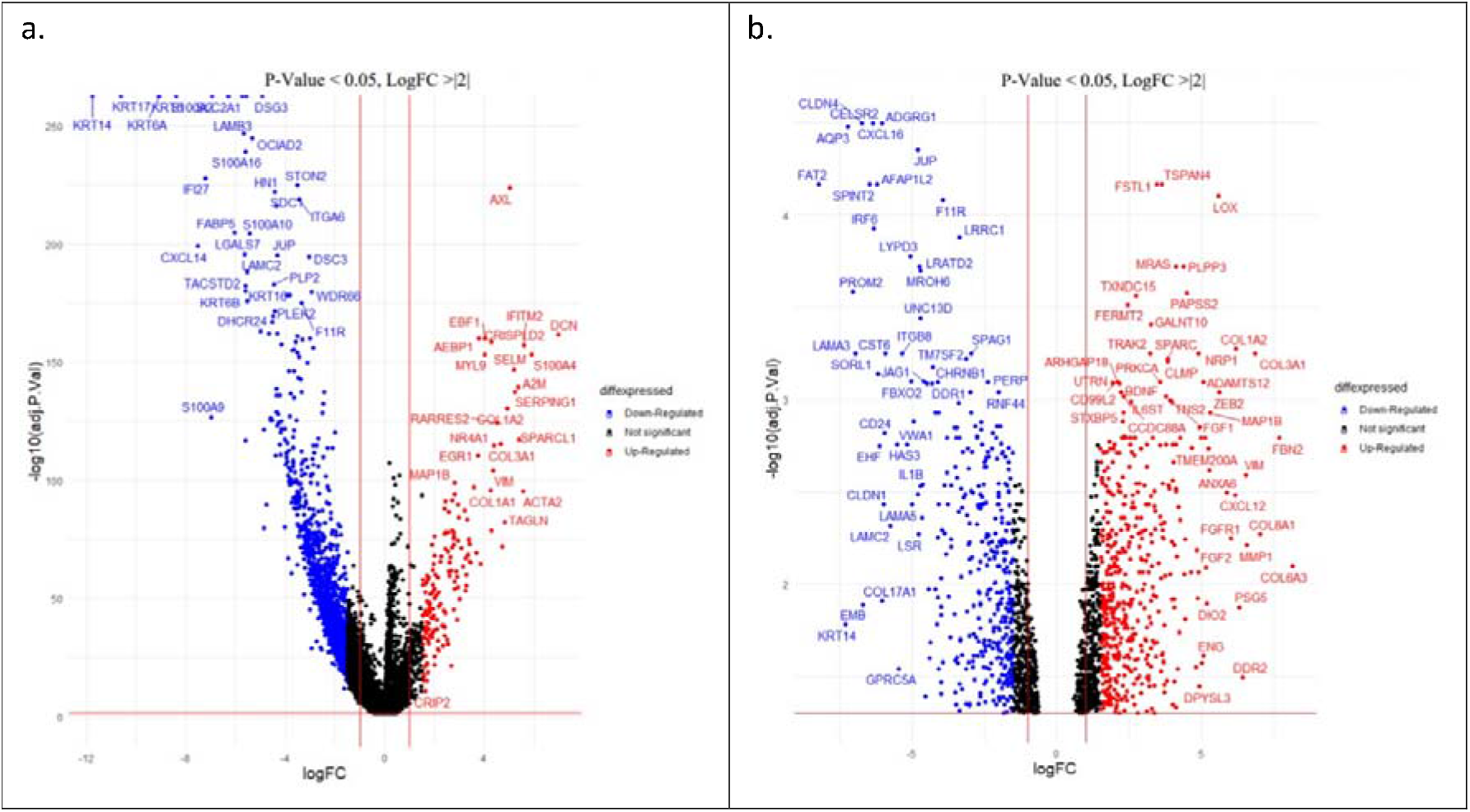
Differentially expressed genes between (a) patient derived cell lines -MhCT08/12-F and MhCT08/12-E; (b) Fibroblast compared to Epithelial from the single cell data GSE103322. Over- and under-expressed genes (logFoldChange > |2| and adjusted P-value <0.05) in fibroblasts are shown as red and blue dots respectively.

### Expression of putative CAF markers in TCGA_HNSC dataset

t-test with RNA-seq counts from dataset GSE135975 indicated only seven (DIO2, MAP1B, COL1A1, SPARC, COL1A2, COL3A1 and TIMP-1) out of the 22 concordant genes to be upregulated in CAFs over normal fibroblasts (Supplementary Table 3). Further using the criterion that a CAF marker should be over-expressed in tumor samples in TCGA Head and Neck cancer dataset, we selected five genes viz. COL1A1, SPARC, COL1A2, COL3A1 and TIMP-1 (Fig. 3).

**Figure 3.**
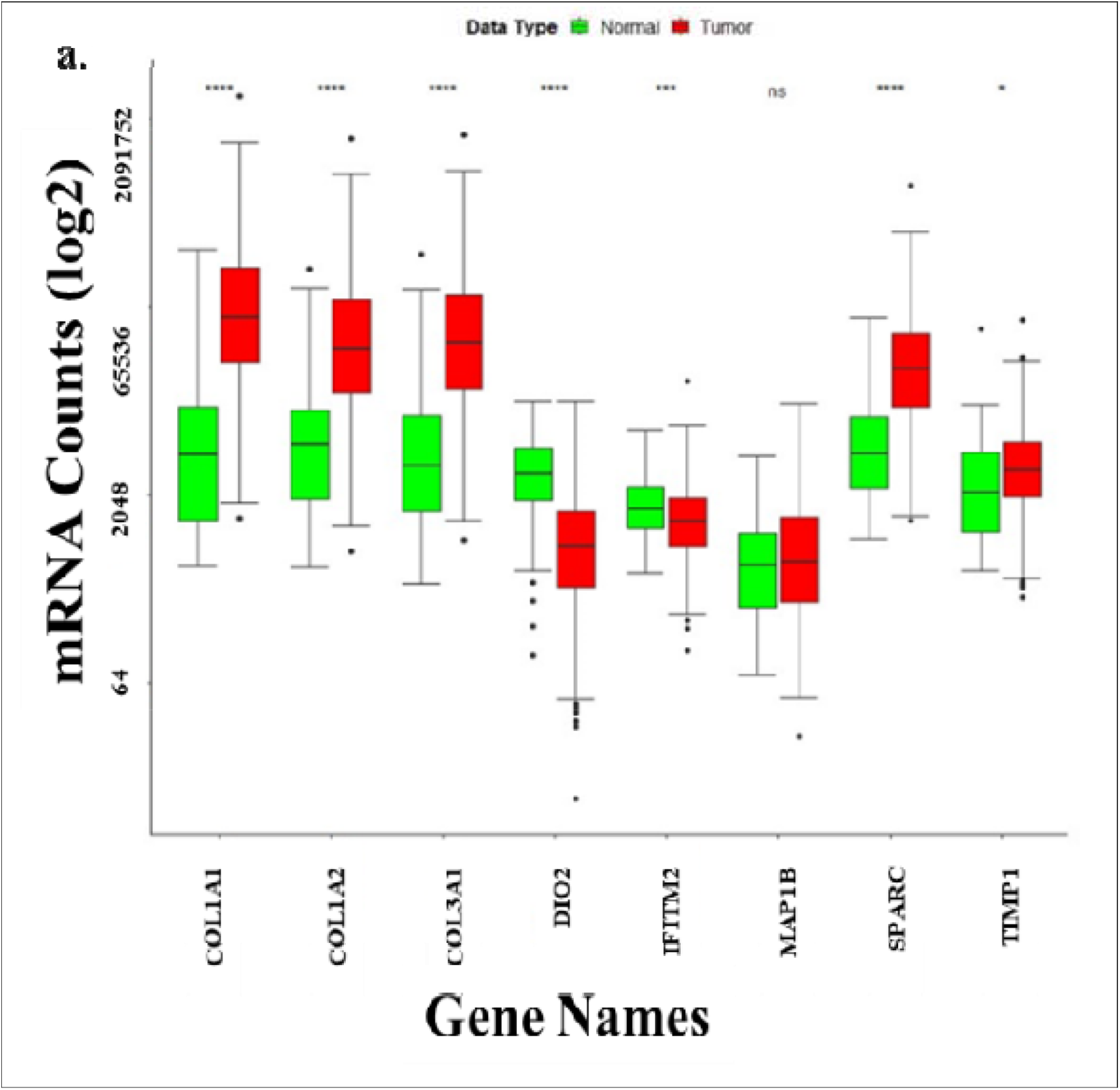
Behaviour of putative CAF markers. mRNA Expression pattern of candidate CAF-marker genes in TCGA HNSC Data. (* P-value <0.05; ns – non significant)

## Discussion

This study presents a data science driven unbiased genome-wide search resulting in selection of CAF markers by differentially expressed genes by comparing the expression of ~400 genes between tumor fibroblast and epithelia and normal fibroblast, in publicly available single cell RNA-seq dataset and in-house patient derived CAF cells of oral cancer.

Earlier used methods for identification of CAF markers majorly constituted of immunofluorescence, immunohistochemistry or flow cytometry based individual pre-selected gene-centric techniques. The novel method described in this study is unbiased towards any particular gene, dealing with a large number of in-house and public domain data obtained from completely independent experiments. This kind of analysis was possible due to the availability of single cell sequencing data (40,57) and a set of novel CAF cells. To our knowledge, the study is the first report of its kind to utilize such a novel platform (58).

CAFs appear to be highly heterogeneous, may be due to the wide variety of potential cellular progenitors from various cellular sites from where they originate (59). Wide heterogeneity of CAFs explain the activation of divergent pro-tumorigenic pathways leading to the expression of varieties of molecular markers

Majority of solid tumors, including head and neck cancer, exhibit SPARC overexpression (60,61). Since SPARC therapy promotes the EMT signalling pathway via AKT activation, exogenous SPARC and tumor-expressing SPARC may be associated to tumor growth in head and neck malignancies (62). An intriguing study by JA Galvan et al. revealed that 87 percent of the patients had SPARC-positive CAFs, which were associated with a number of unfavourable clinicopathological characteristics, such as a higher (y)pT category and lymphatic invasion (63).

On the other hand, human prostate and colon cancer stroma express higher levels of TIMP-1 than healthy stroma aiding in the in vivo progression of both cancer types. TIMP-1 also increases prostate CAF cell proliferation and migration in vitro and promotes ERK1/2 kinase activation in these CAF cells (64). TIMP-1 is said to promote the trans-differentiation of lung fibroblasts into CAFs and decrease apoptosis via SDF-1/CXCR4/PI3K/AKT signalling (65). TIMP-1 has also been associated with the tumor aggressive phenotype, that is, TIMP-1 expression has been shown to decrease in higher grade lesions as compared to well differentiated lesions in oral squamous cell carcinoma (66).

Furthermore, COL family of genes identified in the study, especially COL1A1 and COL1A2 form a part of the transcriptional signature of genes (secreted from CAFs) implicated in promoting the EMT and invasive phenotype in head and neck squamous cell carcinoma (67).

Likewise, EMT gene signature correlated positively with the expression of COL1A2 in colorectal cancer samples (68).

The study is limited by the fact that all the molecules are secretory in nature, hence restricting their usage for sorting CAFs from tumors. However, they can be used for their extensive characterization of origin, even by IHC. In order to characterize the lineage of isolated cells in flow cytometry based experiments, GolgiPlug (cat. no. BD 555029; BD) can be used to arrest the secretion in the established CAF culture to detect the presence of the proposed markers in cells. Curiously, since all the markers identified by the data science approach presented here are secretory in nature indicating that the over expressing CAF-factors reach out more effectively to the tumor as well as the other cells of the microenvironment by spreading through the matrix.

Identification of CAFs utilising the suggested CAF markers opens the door to future selective and therapeutically viable targeting of these cells, enabling the establishment of a successful oral cancer therapy regimen.

## Conclusions

The current study, for the first time shows that COL1A1, SPARC, COL1A2, COL3A1 and TIMP-1 are the most potent CAF markers found and verified at the transcriptome level across multiple CAF types, differentiating CAF from tumour epithelia and normal fibroblast.

## List of abbreviations

ACTA2: Actin Alpha 2
AKT: Alpha Serine/Threonine-Protein Kinase
BSA: Bovine Serum Albumin
CAF: Cancer Associated Fibroblast
CAV1: Caveolin 1
cDNA: Complementary Deoxyribonucleic acid
COL11A: Collagen Type XI Alpha 1 Chain
COL1A1: Collagen Type I Alpha 1 Chain
COL1A2: Collagen Type I Alpha 2 Chain
COL3A1: Collagen Type III Alpha 1 Chain
CRC: Colorectal cancer
CXCR4: C-X-C Motif Chemokine Receptor 4
DAPI: 4ʹ,6-diamidino-2-phenylindole
ddCT: delta-delta Ct
DGE: Differential Gene Expression
DIO2: Iodothyronine Deiodinase 2
ECM: Extracellular Matrix
EMT: Epithelial to Mesenchymal Transition
EpCAM: Epithelial Cellular Adhesion Molecule
EREG: Epiregulin
FACS: Flourescence Associated Cell Sorting
FAP: Fibroblast-activation protein
FBS: Fetal Bovine Serum
FGF7: Fibroblast Growth Factor 7
FSP1: Fibroblast-Specific Protein-1
HGF: Hepatocyte growth factor
HNSCC: Head and Neck Squamous Cell Carcinoma
iCAFs: Inflammatory CAFs
IL-1β: Interleukin 1 Beta
ITGA11: Integrin Subunit Alpha 11
MAP1B: Microtubule Associated Protein 1B
MFAP5: Microfibril Associated Protein 5
MMP11: Matrix Metallopeptidase 11
myCAFs: myofibroblastic CAFs
NCBI: National Center for Biotechnology Information.
NOX4: NADPH oxidase 4
OSCC: Oral Squamous Cell Carcinoma
PBS: Phosphate Buffer Saline
PDAC: Pancreatic Ductal Adenocarcinoma
PDGFRα: Platelet Derived Growth Factor Receptor Alpha
PDGFRβ: Platelet Derived Growth Factor Receptor Beta
PDPN: Podoplanin
PI3K: Phosphatidylinositol-4,5-Bisphosphate 3-Kinase
PLEKHA3: Pleckstrin Homology Domain Containing A3
POSTN: Periostin
qPCR: Quantitative Polymerase Chain Reaction
RIC8B: Resistance To Inhibitors Of Cholinesterase 8 Homolog B
RNA: Ribonucleic acid
ROS: Reactive Oxygen Species
SDF-1: Stromal cell-derived factor-1
SLCC: Stem Like Cancer Cells
SMA: Smooth Muscle Actin
SMTN: Smoothelin
SPARC: Secreted Protein Acidic And Cysteine Rich
STRING: Search Tool for the Retrieval of Interacting Genes/Proteins
TAGLN: Transgelin
TCGA: The Cancer Genome Atlas
TGF: Transforming Growth Factor
TIMP-1: Tissue Inhibitor of Metalloproteinases 1
TME: Tumor Microenvironment
TYW5: TRNA-YW Synthesizing Protein 5
VIM: Vimentin

## Declarations

### Ethics approval and consent to participate

The present study was approved [NHH/MEC-CL-EA-2015-405(A)] by the ethics committee of Narayana Health City (Bangalore, India). Informed consent for the study was obtained from all patients.

### Consent for publication

Not applicable

### Availability of data and materials

The datasets used and/or analysed during the current study are available from the corresponding author on reasonable request.

### Author’s contributions

MD and SKD conceived and designed the study. MD and ND wrote the manuscript. MD, ND, NS and SKD analysed and interpreted the data from public databases and in-house RNA sequencing. MD and SKD reviewed the manuscript. All authors read and approved the final manuscript.

## Acknowledgements

The authors would like to thank Dr. Amritha Suresh for sharing the cell lines used in the study.

