## Supplementary Tables for "Novel CAF-identifiers via transcriptomic analysis in oral cancer patients"

Supplementary Table 1: CAF markers in oral cancer

| S.No | Marker | Techniques used | Drawbacks of the marker |
| --- | --- | --- | --- |
| 1. | Vimentin | IHC (1–7), Microscopy (3,8–10), Western blotting (9–15), Immunofluorescence (10–12,14,16–18), qRT-PCR (5,16,18),  Immunocytochemistry (15), Single Cell Sequencing (19) | Vimentin has ubiquitous expression in the whole fibroblast population as well as multiple other cell types, including macrophages, adipocytes, and the cells undergoing EMT severely limits its utility as a CAF-specific marker (20–22). |
| 2. | ACTA2 | IHC (3–7,23–29), Microscopy (3,9), Western blotting (9,11,12,14,15,25,30–32), Immunofluorescence (11,14,16–18,23,24,31,33–35), qRT-PCR (5,16,18,24,28,30,31), Immunocytochemistry (15,36), Single Cell Sequencing (19), Flow Cytometry (25) | ACTA2 displays fluctuating expression amongst various CAF subtypes (37,38). Additionally, the intracellular location of ACTA2 makes it unsuitable for flow-sorting CAF populations. |
| 3. | FAP | Western blotting (12,30), Immunofluorescence (12,33), qRT-PCR (30), Single Cell Sequencing (19) | Epithelial cells undergoing epithelial-mesenchymal transition (EMT) also exhibit higher levels of FAP (39) |
| 4. | FN1 | IHC (4), Western blotting(12), Immunofluorescence (12), Single Cell Sequencing (19) | Recognizes only myofibroblasts (40) |
| 5. | S100A4 | IHC (4,23), Immunofluorescence (23), qRT-PCR (41), Flow Cytometry (41) | Varies among subtypes as reported in colorectal cancer (38). |
| 6. | THY1 | Immunofluorescence (16), qRT-PCR (16), Single Cell Sequencing (19) | THY1+ fibroblasts are mainly of the reticular lineage (40) |
| 7. | SHH Ligand and GLI1 | IHC, Immunofluorescence (23) | No reports suggesting its absence in epithelial cells, whereas it is present in the epithelial cell development of epidermis, touch dome, hair, sebaceous gland, mammary gland, tooth, nail, gastric epithelium, and intestinal epithelium (42) |
| 8. | COL8A1 and COL11A1 | RNA seq (43) | Not enough experimental validation reports |
| 9. | PDPN, CTGF, JUN, FOS, FGF7, FGF7, TGFBR2/3, MMP11, CAV1, FN1, MMP2, JUNB, IER2, IL8, FOSL1, STAT1, IDO1, ALDH1A1, ALDH3A1, GSTM1, GSTA1, CYP4F11, CYP4F3, ABCC1, GPX2, GSTM2, GSTM3, GSTM4, MYC, TNF, S100A9, PDGFRA, PDGFRL | Single Cell Sequencing (19) | Transgelin (TAGLN) and Periostin (POSTN) (44) in CAFs vary among subtypes, Podoplanin (PDPN), a membrane-bound marker that has been utilised to identify pro-tumorigenic fibroblast subpopulations lacks specificity since it is also expressed in epithelial tumor cells and inflammatory macrophages (45,46). neither PDGFRα nor PDGFRβ are significantly upregulate in CAF populations, due to a more stable expression which is not sensitive to environmental variables like hypoxia, both have been considered as CAF markers in breast cancer (47). |

Supplementary table 2: Techniques for identification of CAF markers: pan-cancer

| S.No | Techniques used | Markers | Cancer type |
| --- | --- | --- | --- |
| 1 | Immunohistochemistry | FAP (48), MFAP5 (49), COL11A1 (50), TN-C (51), PDPN (52), ITGA11 (28), POSTN (53) | Melanocytic skin tumors, Ovarian Cancer, Pancreatic cancer, Prostate cancer, Melanoma, Head and Neck Squamous Cell Carcinoma, Bladder cancer |
| 2 | Immunofluorescence | ACTA2 (54), Vimentin (55), S100A4 (56) | Breast cancer, Human colon tumors, Melanoma |
| 3 | Western blotting | MFAP5 (49) | Ovarian cancer |
| 4 | qRT-PCR | ITGA11 (28) | Head and Neck Squamous Cell Carcinoma |
| 5 | Flow cytometry | PDGFR alpha and Beta (57), S100A4 (56) | Breast cancer |
| 6 | Immunocytochemistry | Vimentin (55) | Human colon tumors |
| 7 | single cell sequencing | ACTA2, FAP, PDPN, CTGF, JUN, FOS, FGF7, VIM, THY1, FGF7, TGFBR2/3, MMP11, CAV1, FN1, MMP2, JUNB, IER2, IL8, FOSL1, STAT1, IDO1, ALDH1A1, ALDH3A1, GSTM1, GSTA1, CYP4F11, CYP4F3, ABCC1, GPX2, GSTM2, GSTM3, GSTM4, MYC, TNF, S100A9, PDGFRA, PDGFRL (19)  BGN, LUM, CCL19, CEBPD, and ID3 (58)  ADAMTSL2, SLCO2A1, CD4, HMGXB3, GCN1, and LUC7L3 (59) | Head and Neck Squamous Cell Carcinoma, Breast Cancer |

Supplementary Table 3. P-Value for the selected marker genes in CAF Versus Fibroblast data using t.test.

| **HGNC** | **Difference between the counts of CAF and Fibroblast** | t-test p-value |
| --- | --- | --- |
| DIO2 | 4270.458 | 0.000443 |
| MAP1B | 42710.75 | 0.001093 |
| IFITM2 | -495.833 | 0.001899 |
| COL1A1 | 127671.2 | 0.008181 |
| SPARC | 60279.13 | 0.015402 |
| COL1A2 | 192046.8 | 0.019645 |
| COL3A1 | 119438.3 | 0.022073 |
| TIMP1 | 10349.42 | 0.023561 |

Supplementary Table 4: List of Reagents used in the study

| S.No | Product | Catalogue Number | Source |
| --- | --- | --- | --- |
| 1 | AMV Reverse Transcriptase | M0277S | NEB |
| 2 | anti-mouse Alexa488 (1:200) | A11029 | Invitrogen |
| 3 | anti-SPARC antibody (used at 0.5mg/mL) | MAB941-SP | R&D systems |
| 4 | anti-TIMP1 antibody (used at 0.5mg/mL) | MAB970-SP | R&D systems |
| 5 | BSA | TC194 | HiMedia |
| 6 | Chloroform | 496189 | Sigma |
| 7 | DAPI | F6057 | HiMedia |
| 8 | EpCAM (working solution) | AN820 | BioGeneX |
| 9 | Ethanol | MB228 | HiMedia |
| 10 | FBS | 10270-106 | Gibco^TM^ |
| 11 | FSP-1 (used at 2mg/mL) | F4771 | Sigma |
| 12 | Golgi plug | 554724 | BD |
| 13 | Isopropanol | DB4DF64078 | Merck |
| 14 | KAPA SyBr green Universal | KK4600 | Kapa |
| 15 | Methanol | AS059 | HiMedia |
| 16 | Nuclease free water | 821739 | MP |
| 17 | PanCK (used at 1:200 dilution) | 4545 | CST |
| 18 | PBS | 10010-023 | Gibco |
| 19 | Penicillin- Streptomycin | 15140122 | Gibco^TM^ |
| 20 | Qubit RNA Assay kit BR | Q10210 | Invitrogen |
| 21 | RPMI- 1640 | AT222A | HiMedia |
| 22 | Triton-X 100 | 10655 | Fischer Scientific |
| 23 | TRIzol | 15596018 | Invitrogen |
